# Does methylation of NFATC1 and C-FOS genes are associated with post-menopausal osteoporosis?

**DOI:** 10.1101/2020.02.19.955682

**Authors:** Rasime Kalkan, Özgür Tosun

**Author notes:** **Corresponding Author:** Assoc. Prof. Rasime Kalkan, Near East University, Faculty of Medicine, Department of Medical Genetics, Cyprus, 99138. **Author Contribution:** RK supervised the project. RK conceived and planned the experiments. OT carried out the statistical analysis. RK. carried out the experiment. RK and OT wrote the manuscript. Both RK and OT. authors contributed to the final version of the manuscript. All authors provided critical feedback and helped shape the research, analysis and manuscript.

## Abstract

Genetic and epigenetic factors have an important role during the development of osteoporosis. RANK/RANKL pathway is important for the bone remodeling and NFATC1 and c-FOS are the downtargets of this pathway. Here, we report methylation status of *NFATC1* and *C-FOS* genes in post and premenopausal cases. In this study 30 pre-menopausal and 35 post-menopausal cases were included. MS-HRM was used for identification of *NFATC1* and *C-FOS* metyhylation.

NFATC1 were methylated in 11 of the 35 post-menopausal women and C-FOS were methylated in 6 of the postmenopausal women (p >0.005). Here, we found statistically significant association between unmethylation of NFATC1 and post-menopausal status. This result explains the epigenetic regulation of osteoclasts during the menopausal transition and our results can be used for epigenetic explanation of post-menopausal osteoporosis for the first time in the literature. Although the limited number of studies in this field makes our results crucial. Therefore, our results showed great value of epigenetic profile of post-menopausal women.

## INTRODUCTION

Nuclear Factor of Activated T-cell gene family contains 5 members; which are NFATC1, NFATC2, NFATC3, NFATC4 and NFATC5 ^1^ and they regulated by the calcium signaling pathway ^1^. This pathway plays an important role during the regulation of the different systems, for instance; immune system ^2^, skeletal system and circulatory system ^2,3,4^. NFATc1 regulates osteoclast specific genes to be able to regulate osteoclast differentiation, like TRAP, cathepsin K, calcitonin receptor and c-Fos ^5,6,7,8^. Osteoclast differentiation triggers NFATc1 expression and this pathway was under control of RANKL [Receptor activator of nuclear factor-kappa B (NF-κB) ligand), NF-κB and c-Fos signaling ^9^. RANK/RANKL is the important pathway for the bone remodeling ^10^. RANKL-RANK activates c-Fos, which was triggers NFATC1 expression ^8^. The interaction between C-fos and osteoclast differentiation observed in mice and researchers showed the mice whose c-fos knockout they developed osteopetrosis ^5^, ^11^.

Due to the genetic and epigenetic factors, osteoporosis reported as a multifactorial disease ^12^. Several candidate gene association studies had been done but the etiology and molecular mechanism of disease is not clearly explained. During the menopause, the deficiency of estrogen causes increased level of FSH which accelerates bone loss ^13^. Differential gene expression of bone cells has been done to solve this complex interaction ^14,15^. Liu and colleagues used circulating monocytes for genome wide differential gene expression analysis in osteoporotic and non-osteoporotic postmenopausal Caucasian females and they identified differential gene expression in m circulating monocytes ^16^. In general studies analyzed the effect of estrogens, corticosteroids and oxidative stress on osteoblast cells ^15,17^ and showed the gene expression profile or polymorphisms of premenopausal and primary osteoporosis cases ^15,18^.

Based on our knowledge, there are no epigenetic studies of NFATC1 and c-FOS in post-menopausal women. In this study, we aim to analyze methylation status of NFATC1 and C-FOS genes in post and premenopausal cases and to show their association between menopausal osteoporosis. Based on the current literature, this is the first epigenetic study on postmenopausal women which shows the post-menopausal osteoporosis and NFATC1 interaction.

## MATERIALS AND METHODS

The study protocol was approved by the Local Ethics Committee (YDU/2016/42-352). Written informed consent was obtained from all participants.

Genomic DNA was extracted from blood samples according to the Qiagane AllPrep DNA/RNA/Protein isolation kit and its quantity was measured using a NanoDrop ND-1000 Spectrophotometer (Thermo Fisher Scientific).

### Subjects

In this study, cases were divided into two groups. One group consist of 35 post-menopausal cases which their mean age was 56.7± 4.9 and contains 30 pre-menopausal cases and their mean age was 33.5 ± 6.9. All of the participants were lack of the hypertension, liver, kidney, diabetes, thyroid or cardiovascular disease and medical therapy history had been questioned. They were neither received any medications nor participated in any dietary or exercise program during the study. All participants were genetically unrelated postmenopausal females. Subjects with un-natural menopause, took medications such as anxiolytics, anti-depressants, exogenous hormone, women who have serious disease or mental retardation, smoking, alcohol usage and have a weight loss therapy, food allergies, heart disease history, insulin-dependent diabetes, type 2 diabetes, kidney disease or liver disease were excluded. Individuals fitting to menopause for > 1 year were recruited.

### Determination of NFATC1 and c-FOS methylation status

Bisulfite modification was applied according to the manufacturers’ protocol (EpiTect Bisulfite Kit (Qiagen, Manchester, UK). and 1.3 μg DNA was used for bisulfite treatment reaction. QIAGEN Rotor Gene Q used for MS-HRM (Methylation Sensitive High Resolution Melting) analysis to detect the methylation status of *NFATC1* and *c-FOS* genes and also universal methylated and unmethylated DNA (EpiTect Control DNA Set, Cat No./ID: 59568) were used as a control in each reaction. The primers for each were designed according to the EpiTect® HRM™ PCR Handbook^19^.

### Statistical analysis

The statistical analyses and their associations with patient characteristics were performed by Pearson’s chi-squared test and two tailed Fisher’s exact test. Calculations were performed using SPSS 16.0 software (SPSS Demo for Mac, Chicago, IL, USA), with a statistical significance of p< 0.05.

## RESULTS

The mean age of 30 pre-menopause patients was 33.5 years (mean ± Std. Deviation, 33.5 ± 6.9) and 56.7 (mean ± Std. Deviation, 56.7± 4.9) was the post-menopause patients. There were no significant differences between the premenopausal and postmenopausal group in BMD (Bone mineral density) (p>0.005).

NFATC1 promotor methylation was detected in 11 of the 35 post-menopausal women (31.4%) (Figure 1) and unmethyalted in 24 of the 35 post-menopausal women (68.8%) (Figure 2).

**Figure 1:**
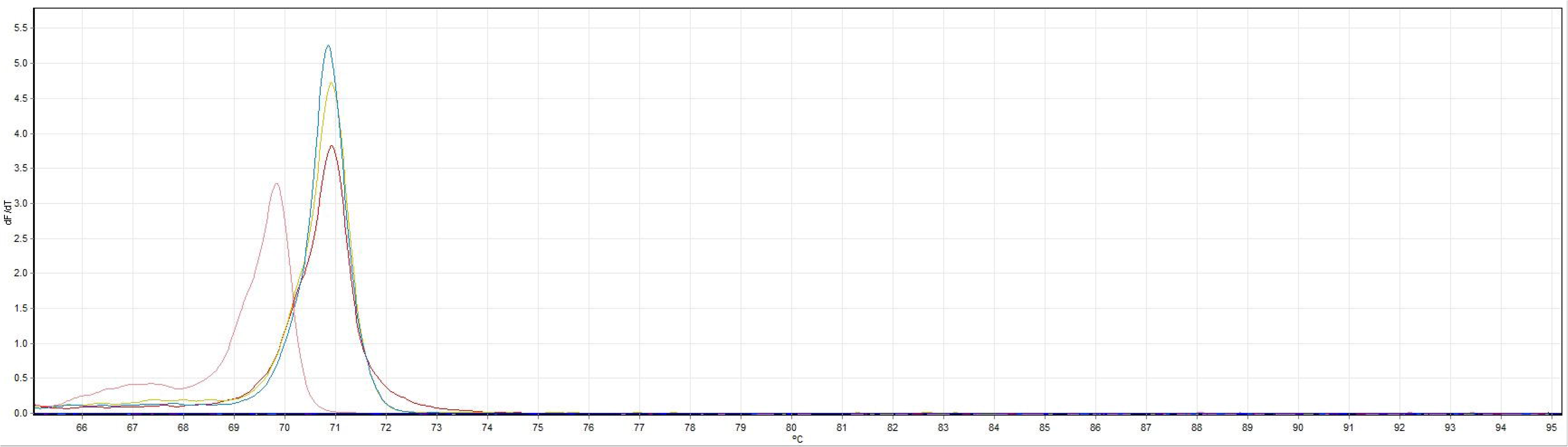
Methylated NFATC1 patients. NFATC1 unmethylated control was orange, methylated control was red. Patient number 41, 34 were methylated.

**Figure 2:**
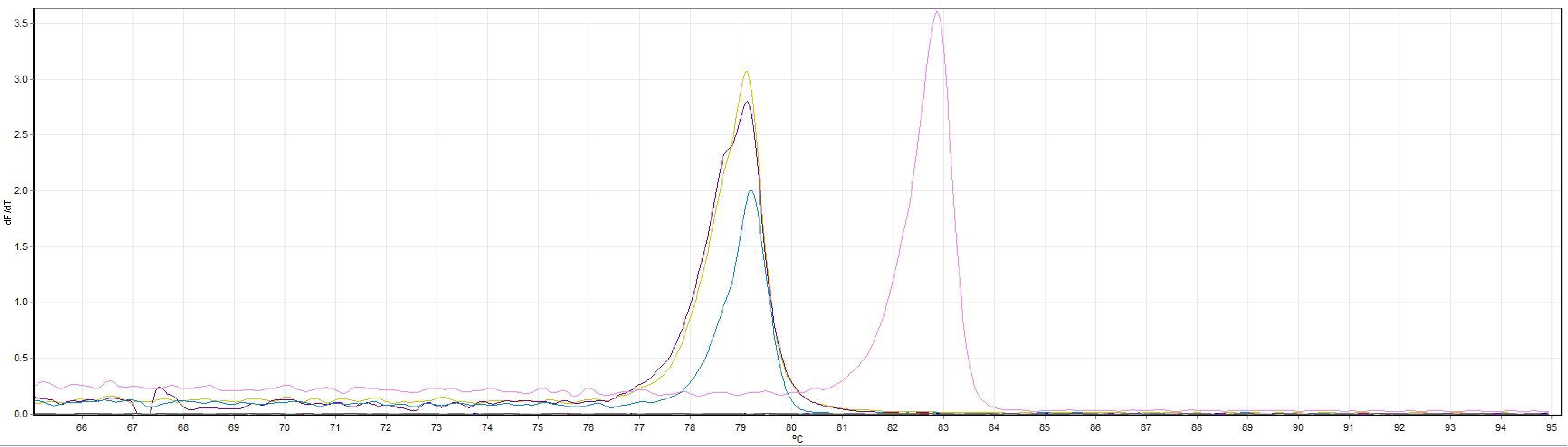
Unmethylated NFATC1 patients. NFATC1 unmethylated control was yellow, methylated control was purple. Patient number 1, 15 were unmethylated.

NFATC1 promotor methylation was detected in 19 of the 30 (63.3%) control sample (Pre-menopausal cases) and unmethylated in 11 of the 30 control sample (36.7%). Here, we found statistically significant correlation between post-menopause and unmethylation of NFATC1 promotor (p = 0,010). C-FOS promotor methylation was detected in 6 of the 35 (17.1%) post-menopausal women (Figure 3) and unmethylated in 29 of the 35 post-menopausal women (82.9%) (p >0.005).

**Figure 3:**
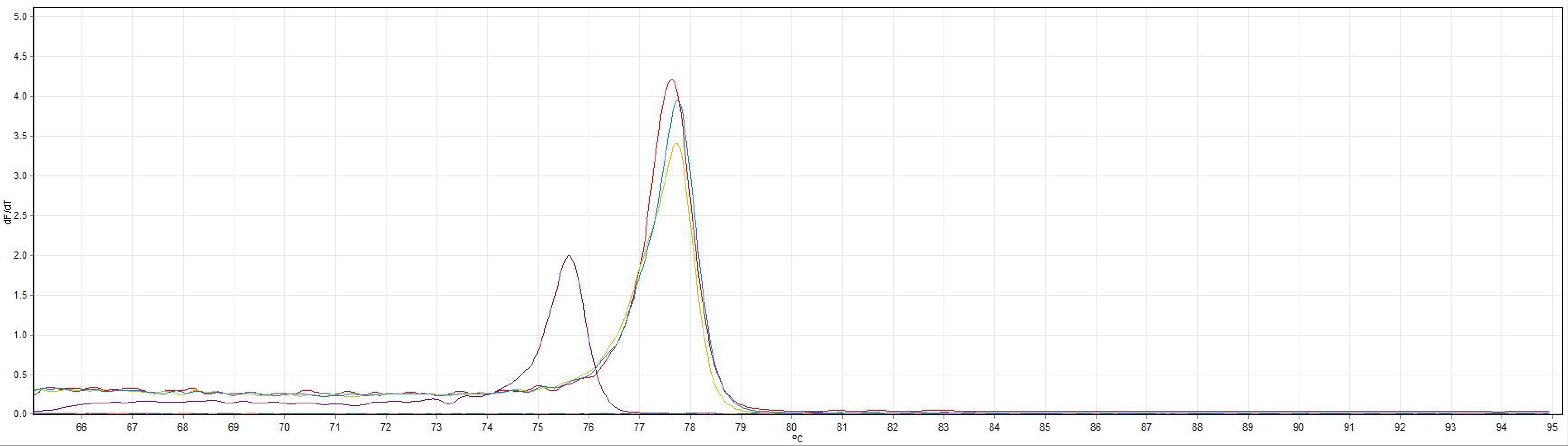
Methylated c-FOS patients. C-FOS unmethylated control was purple, methylated control was red. Patient number 8, 19 were methylated.

C-FOS promotor methylation was detected in 5 of the 30 (16.7%) control sample (Pre-menopausal cases) and unmethylated in 25 of the 30-control sample (83.3%). Besides that, there were no statistically significant association identified between menopause and methylation status (p>0.05) (Table 1).

## DISCUSSION

RAS signaling pathway is important for the osteoclast survival and triggers downstream gene activations ^20^. NFATC1 and c-FOS are the downtargets of RAS pathway which they have an important role during the osteoclastogenesis ^20^. NFATc1 regulates osteoclast specific genes during the regulation of osteoclast differentiation and c-FOS is one of the genes which NFATC1 triggers their expression ^5,6,7,8^. Raggatt and colleagues demonstrated that NFATC-1 is one of the important transcription factor during the differentiation of osteoclast precursors ^21^.

Decreased level of estrogen triggers several physiological and hormonal problems and postmenopausal osteoporosis is one of the consequences of decreased level of estrogen ^22,23^. Gavali and colleagues demonstrated the effect of estrogen on osteoclast differentiation and they showed lack of the estrogen causes failure of apoptosis and inhibits activation of NFATC1 and c-Src ^24^. Kim and colleagues tested the extract of Lycii Radicis Cortex (LRC) on an ovariectomized (OVX) rat model and they showed suppression of RANKL-induced osteoclast differentiation and also demonstrated inhibitory effect of LRC on NFATC1 expression ^25^. On the other hand, Kim and colleagues showed the overexpression of NFATC1 induced differentiation of osteoclasts ^26^. Also, Zhao and colleagues showed that NFATc1 is an important transcription factor for RANKL-mediated osteoclast differentiation^27^ and Winslow and colleagues demonstrated lack of the osteoclast differentiation and osteoporosis was observed on NFATc1 knock-out mice ^28^. Researchers also showed the inhibition of NFATC1 had an important role during the RANKL-induced osteoclast differentiation ^29^.

Many studies have shown the effects of inhibition of NFATC1 and c-FOS on osteoclasts and postmenopausal osteoporosis ^8,20,29^. As we mentioned previously c-Fos is another member of RANK/RANKL pathway and stimulation of c-Fos causes the activation of NFATC1 ^11,27^. The effects of the c-Fos deletion has been shown in mice which causes defective osteoclast differentiation and osteopetrosis and Kim and colleagues showed inhibitory effects of LRC exerted on c-Fos mRNA ^25^.

The interaction between NFATC1 and C-FOS has been studied in many years with animal models in *in vitro* conditions. Here we studied NFATC1 and C-FOS methylation status in blood samples of post and pre-menopausal cases. In this study, we showed statistically significant association between post-menopause and unmethylation of NFATC1 promotor (p = 0,010) and methylation of NFATC1 and pre-menopause. NFATC1 is important for the osteoclast differentiation, which means important for the bone resorption. According to our results, osteoporosis does not expected condition in pre-menopausal cases and methylation data also confirms this condition and in post-menopausal cases and the unmethylation of NFATC1 promotor (p = 0,010) means the differentiation of osteoclasts were induced. These results could be used to epigenetic explanation of the post-menopausal osteoporosis. Also, future studies should be design to inactivation of NFATC1 pathway as a possible targeted therapy strategy in post-menopausal osteoporosis. On the other hand, we did not find any significant associations between menopause and C-FOS methylation (p>0.05). As far we know epigenetic alterations are tissue specific but we hope our results should be shed light to the further studies in this field and triggers researchers to used bone and related tissues for methylation analysis of this pathway. Until know, there are no epigenetics study in this field and we can conclude that NFATC1 gene was hypermethylated during the pre-menopausal term and it could be unmethylated during the post-menopausal term. We hope our results will led epigenetic studies in this field.

### Conclusions

In conclusion, we found no significant association between the C-FOS gene methylation and menopause. But we found significant association of unmethylation of NFATC1 and post-menopausal status. NFATC1 and osteoclast formation interaction has been shown by the in vitro studies. In this study, we can conclude that the epigenetic silencing of the *NFATC1* gene can be raise osteoclast activity and it is directly related with post-menopasusal osteoporosis. These results will shed light further epigenetic studies in postmenopausal cases.

This study is the first epigenetic study that investigated the NFATC1 and FOS methylation in post-menopausal cases and shows the interaction between epigenetics and post-menopausal osteoporosis. Although the limited number of sample size in our study and lack of epigenetic studies in this field proves our results crucial and therefore, our results showed magnitude of epigenetic profile of Turkish Cypriot post-menopausal women.

Future studies in larger samples of postmenopausal women focused in the study of the different gene methylations and this will help to clarify potential effects of gene methylation in menopause.

Table 1. Methylation status of NFATC1 and c-FOS in post and pre-menopausal cases.

## Compliance with Ethical Standards

The authors declare that they have no conflict of interest.

## Acknowledgments

This study was supported by Near East University Scientific Research Project Unit-Grant Number: SAG-2016-2-012. We would like to thank all the participants involved in this study.

