## Supplemental Table 1 for "Does methylation of NFATC1 and C-FOS genes are associated with post-menopausal osteoporosis?"

Table 1. Methylation status of NFATC1 and c-FOS in post and pre-menopausal cases.

|  | Post-Menopausal | Pre- Menopausal | P Value |
| --- | --- | --- | --- |
| **NFATC1 Methylated** | 11(%31.4) | 19(%63.3) | P=0.010 |
| **NFATC1 Unmethylated** | 24 (%68.8) | 11 (%36.7) |  |
| **C-FOS Methylated** | 6(%17.1) | 5(%16.7) | P>0.05 |
| **C-FOS Unmethylated** | 29(%82.9) | 25(%83.3) |  |
